# Inter-generational Consequences for Growing C.elegans in Liquid

**DOI:** 10.1101/467837

**Authors:** Itamar Lev, Roberta Bril, Yunan Liu, Lucila Inés Ceré, Oded Rechavi

## Abstract

In recent years, studies in *Caenorhabditis elegans* nematodes have shown that different stresses can generate multigenerational changes. Here we show that worms that grow in liquid media, and also their plate-grown progeny, are different from worms whose ancestors were grown on plates. It has been suggested that *C.elegans* might encounter liquid environments in nature, although actual observations in the wild are few and far between. In contrast, in the lab, growing worms in liquid is commonplace, and often used as an alternative to growing worms on agar plates, to control the composition of the worms’ diet, to starve (and synchronize) worms, or to grow large populations for biochemical assays. We found that plate-grown descendants of M9 liquid media-grown worms were longer than control worms, and the heritable effects were apparent already very early in development. We tested for the involvement of different known epigenetic inheritance mechanisms, but could not find a single mutant in which these intergenerational effects are canceled. While we found that growing in liquid always leads to inter-generational changes in the worms’ size, trans-generational effects were found to be variable, and in some cases the effects were gone after 1 -2 generations. These results demonstrate that standard cultivation conditions in early life can dramatically change the worms’ physiology in adulthood, and can also affect the next generations.

## Introduction

“*You could not step twice into the same river* (Heraclitus)

In comparison to Mendelian inheritance, which entails faithful transmission of discreet packets of information which do not blend or dilute [1], heritable epigenetic data is “soft”, or “fluid”, as it changes as a function of time, and in response to physiological processes [2].

Generally, mechanisms which allow trans-generational epigenetic effects, namely transmission of information to progeny not exposed to the original trigger, are still poorly understood [3]. However, in *C.elegans* substantial understanding has been gained regarding the molecular mechanisms that allow transmission of responses via small RNAs across multiple generations (Reviewed in: [4]). Further, many studies with *C.elegans* have demonstrated that different environmental challenges leave a trace in the progeny, for example viral infections [5–7], starvation [8–11], high temperatures [12–14], high osmolality [15,16], and exposure to toxins [15]. Such stressors can lead to short or long-term gene expression alternations and physiological changes. The inheritance of some of these responses is associated with changes in heritable small RNAs [8,13,14] and/or is correlated with alternation in histone modifications (for example, of H3K4me [15,17].

In the lab, *C. elegans* are commonly grown on agar plates, but when large numbers of worms are needed, worms are often grown in liquid media [18]. *C.elegans* are grown in liquid for other purposes as well, for example to synchronize the worms’ growth at the L1 stage [18], to control for the composition of the worms’ diet [19], and sometimes for studying starvation [18]. Liquid media was also used to study how swim exercise affects the worm’s physiology [20]. In addition to *C. elegans*, other nematodes, such as the entomopathogenic species *Steinernema* [21,22] and *Heterorhabditis* [23,24] are routinely grown in liquid cultures, and recently, growth in liquid medium of different *Pristionchus* species was shown to generate striking phenotypic differences [25] (see also discussion).

In nature, *C. elegans* worms are most often isolated from solid decomposing plant material. However, there are rare reports of isolation of *C. elegans* from freshwater as well [26]. In addition, it has been hypothesized, but not yet documented, that *C. elegans* could encounter liquid environments in the wild, in cases of excessive decomposition, especially of fruits [27].

Here, we report that growth of worms in liquid media changes the morphology of the liquid-grown worms and also the morphology of their plate-grown progeny. We found that plate-grown descendants of liquid-grown worms are on average longer than control worms. This morphological change is apparent early during the worms’ life, and persists even in spite of experimental synchronization of the worms’ development. In addition, the inheritance of this phenotype does not appear to depend on pheromone production or any single known epigenetic inheritance pathway, as we could not find a single mutant in which these inter-generational effects would be canceled. While the effect is robustly inherited to the immediate F1 progeny, the inheritance to the F2 and the F3 generations exhibits variability. These results demonstrate that commonplace cultivation conditions can dramatically change the worms’ physiology in adulthood, and can also affect the next generations. We suggest that the growth and handling conditions of at least a few generations should be monitored before experimentation begins.

## Results

Even in the lab, *C. elegans* often encounters different environments, for example, worms are frequently starved or grown in slightly stressful temperatures, either accidentally or intentionally. It is very common to synchronize or control the speed of the worms’ growth, by starving them or by switching the growth temperature. Similarly, in many cases, researchers are forced to grow temperature-sensitive worms in low temperatures, or to cultivate the worms in high temperatures to avoid transgene silencing [28]. Contaminations with different pathogens are very common in *C.elegans* cultures, and while this has not been studied in detail, such infections could also leave heritable imprints [29]. As described above, worms are often grown in liquid in the lab (the standard liquid used is M9 buffer,[18]). In the course of conducting many experiments with liquid-grown worms, we noticed that after growth in liquid medium, the worms exhibit a different morphology, in comparison to plate-grown worms. We first set out to characterize this phenomenon, and to examine whether the effects could be inherited (**Figure 1A**).

**Figure 1.**
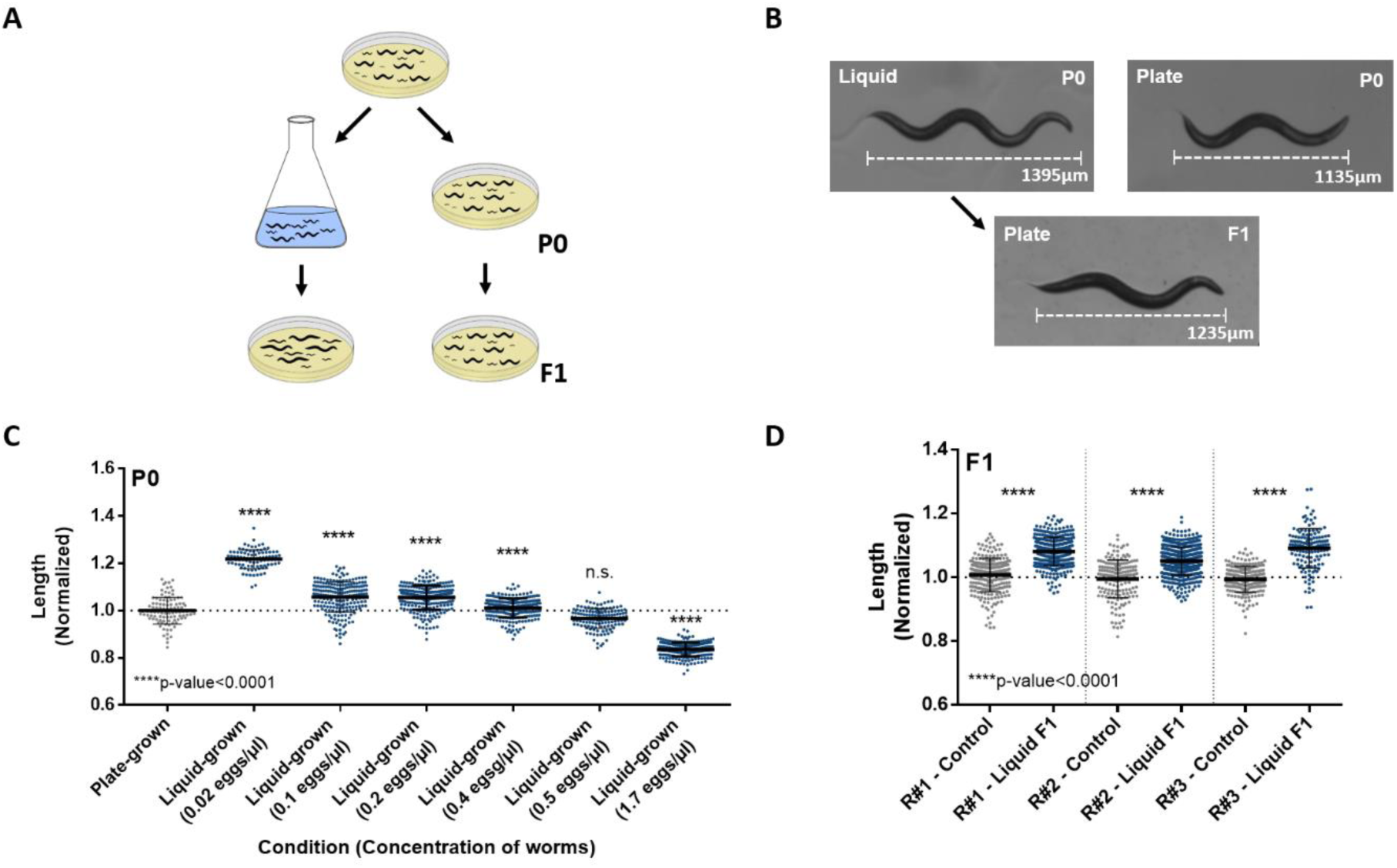
Progeny of liquid-grown worms have longer morphology. A. Experimental scheme of inter-generational inheritance of growth in liquid medium. Eggs were produced from Isogenic worm lines and placed either on plates or in liquid medium. When the parental P0 worms reached adulthood, they were placed on plates to lay their eggs. When the resulting F1 progeny reached adulthood, their body length was measured.
B. Representative images of adult plate-grown (P0), liquid-grown (P0) or plate-grown progeny (F1) of liquid-grown worms. The differences in length between the photographed worms match the average length differences found between the examined populations. A scale bar that shows the total *stretched* length of the worm is presented.
C. Liquid-grown worms are longer when grown in low concentration. The normalized length of the parental plate-grown, and liquid-grown worms are shown. Worms were grown in liquid in different concentrations (x-axis). The measured length was normalized to the plate-grown worms. ^∗∗∗∗^p-value<0.0001, One-way ANOVA, Sidak’s multiple comparisons test.
D. Plate-grown progeny of worms grown in liquid medium are longer. The length of F1 plate-grown progeny of liquid-grown worms and progeny of plate-grown controls are shown. The measured length was normalized to the corresponding control worms. Data from three independent biological repeats are presented (N>150 worms per group). The p-value was determined using Two-way ANOVA with the two categorical variables: A. biological condition (growth in liquid or on plates) and B. biological repeat (independent experiments conducted on separate groups of animals, on different days). Both factors have a significant effect on the measured worm length (p-value <0.0001). In addition, an interaction effect was found as well (p-value <0.0001). The biological condition explained far more variance than the biological repeat or the interaction (32.44% compared to 2.07% or 1.69%, respectively).

### Progeny of liquid-grown worms are longer

We noticed that worms that grow in liquid become narrow, and that their body length becomes more variable in comparison to plate-grown worms. These observations are in accordance with a previous observation [18], and a recent study showing that worms grown in axenic (chemically defined and bacteria-free) liquid medium (specifically the “Habitation and Reproduction” medium used by NASA when growing worms in space [19]) are also narrower and longer than plate-grown worms [30].

We found that the worms’ concentration in liquid affects their size (p<0.0001, One-Way ANOVA), and that while in low concentrations the liquid-grown worms are longer (**Figure 1B and C**), in very high concentrations, liquid-grown worms become shorter (perhaps in this condition not enough food is available). As starvation is known to generate heritable effects in *C.elegans* [8–11], in all the subsequent experiments we grew worms in liquid in low concentrations (maximum 0.05 worms per microliter, see methods). In this concentration, liquid-grown worms are ~22% longer than plate-grown worms (p<0.0001, Sidak’s multiple comparisons test, **Figure 1B and C**).

Next, we tested whether plate-grown progeny of liquid-grown worms differ in their body length (**Figure 1A and B**). Interestingly, we found that the F1 progeny of liquid-grown worms were on average ~8% longer than control worms (p<0.0001, Two-Way ANOVA, **Figure 1B and D**). The heritable effect was very robust, repeated across many biological repeats, and was independent of the concentration of the parent worms in the liquid medium (**Figure S1**, in addition, experiments were replicated by multiple different experimenters).

When placed in liquid media worms change their movement, to a swim-like behavior, which involves continuous “thrashing”. Previously this swim-like behavior was used to study the effects of exercise on the worm’s physiology [20]. We wondered whether this energetically demanding behavior could be involved in the morphological change found in the liquid-grown worms and their plate-grown progeny. To test this we examined the morphology of liquid-grown immobile *unc*-*119* mutants across generations. We found that liquid-grown *unc*-*119* mutants do not become longer in liquid (p<0.0001, Two-Way ANOVA, **Figure S2A and B**). While experimentation with such *Unc* mutants is challenging (the mutants exhibit higher length variability, and do not grow well in liquid), the plate-grown progeny of liquid-grown mutants were nevertheless longer than controls (p<0.0001, Two-Way ANOVA, **Figure S2C**). These results suggest that swimming in liquid is not required for the intergenerational effect.

Recently, Celen et al analyzed the transcriptome of worms that grew in a special axenic liquid medium [31]. They found that the genes that were up-regulated in axenic media were enriched with genes related to cuticle formation and morphology. Further, some transcriptional changes were also inherited to plate-grown F1 progeny. The authors did not examine whether the morphology of the progeny of the worms that grew in axenic medium changed, however it is possible that these transcriptional changes correspond to the inherited morphological changes that we observe in the progeny of M9 liquid grown worms.

### The change in the morphology of the progeny of liquid-grown worms is apparent early during development, and persists even when the progeny’s development is synchronized

It is possible that the progeny of the liquid-grown worms is longer, since these worms develop faster. To test this possibility, we monitored the worms’ length throughout development. To our surprise, we found that progeny of liquid-grown worms were longer already as eggs (p<0.0001, Two-Way ANOVA, **Figures 2A**). The progeny of the liquid-grown worms were also longer immediately after hatching, at the L1 stage (**Figures 2B and S3**, p=0.022, see statistical methods), maintained the elongated morphology throughout development, and their rate of development was identical to the control group (**Figures 2B and S3**, p=0.7806, see statistical methods). Moreover, the effect persisted in adults, even when we synchronized the worms’ development at the L1 stage by starvation (for 12 hours, see methods. p<0.0001, Two-way ANOVA, **Figure 2C**). In summary, the elongated morphology of the F1 progeny of liquid-grown worms does not stem from differences in the rate of development.

**Figure 2.**
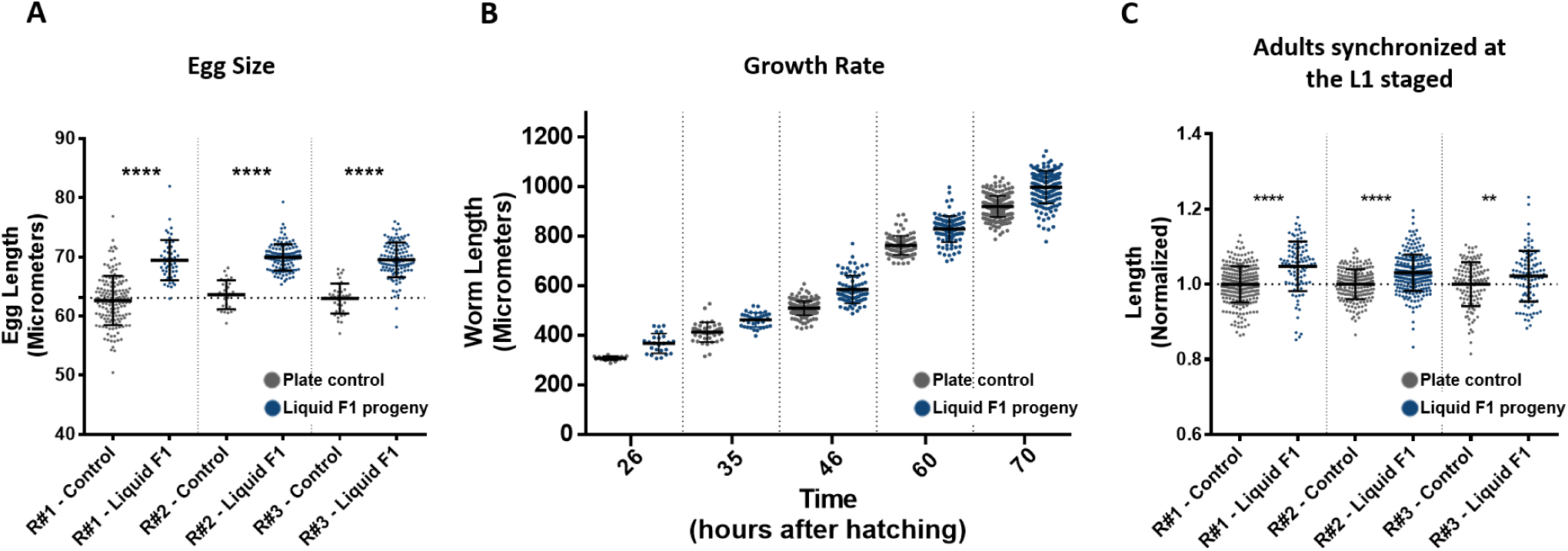
Progeny of liquid-grown worms are longer already early in development. A. The length of eggs laid by liquid-grown worms and plate control worms. Data from three independent biological repeats are presented (N>30). ^∗∗∗∗^p-value<0.0001, Two-way ANOVA (biological condition is significant p<0.0001, biological repeat and interaction effect factors are not significant: p= 0.8642 and p=0.2045, respectively). Sidak’s multiple comparisons test p-value is presented.
B. The length of progeny of liquid-grown worms and control worms are presented at different stages of the worm’s development.
C. The length of adult worm progeny of liquid-grown and plate-grown controls, 72 hours after being released from starvation-induced developmental arrest at the L1 stage. Data from three independent biological repeats are presented (N>100). ^∗∗∗∗^p-value<0.0001, ^∗∗^p-value=0.0066. Two-Way ANOVA (biological condition is significant p<0.0001, biological repeat and interaction effect factors are also significant p=0.0093. These factors explain 0.81% of the variance, while the biological condition explain 7.96% of the variance). Sidak’s multiple comparisons test p-value is presented. Error bars represent standard deviations.

### Progeny of liquid-grown worms are longer even in mutants defective in chemosensation and pheromone biosynthesis

In response to various environmental changes, *C. elegans* alter the composition of the molecular social cues (or pheromones) that they secrete [32,33]. In turn, the secreted pheromones greatly affect the worms’ physiology [34,35]. For example, in response to crowded culture conditions, *C. elegans* secrete pheromones which cause starved larval worms to arrest their development and enter the dauer stage (an alternative developmental stage during which worms can endure harsh conditions) [35]. We examined whether sensation of pheromones or other secreted molecules could be the factor that transforms the progeny of the liquid grown worms. To test this, we conducted experiments with mutants defective in pheromone production in general (*daf*-*22*) and dauer pheromones sensation specifically (*srg*-*36/37*). We also examined mutants that are generally defective in chemosensation (*che*-*2*).

DAF-22 is a fatty acid beta oxidase important for the production of short-chain ascarosides [36] and dauer pheromones [37]. SRG-36/37 are two redundant G-protein-coupled receptors for the dauer pheromone ascaroside C3 [38]. Mutations in these genes, which confer resistance to dauer formation, were identified in two independent long-term liquid-grown strains (in worms that were grown in liquid for many decades, [38]). We found that liquid-grown *daf*-*22* and *srg*-*36/37* mutants generate a functional intergenerational response and similarly to wild type animals, plate-grown progeny of the liquid-grown mutants were longer than controls (**Figure 3**). Thus, pheromones are probably not involved in this inter-generational effect.

**Figure 3.**
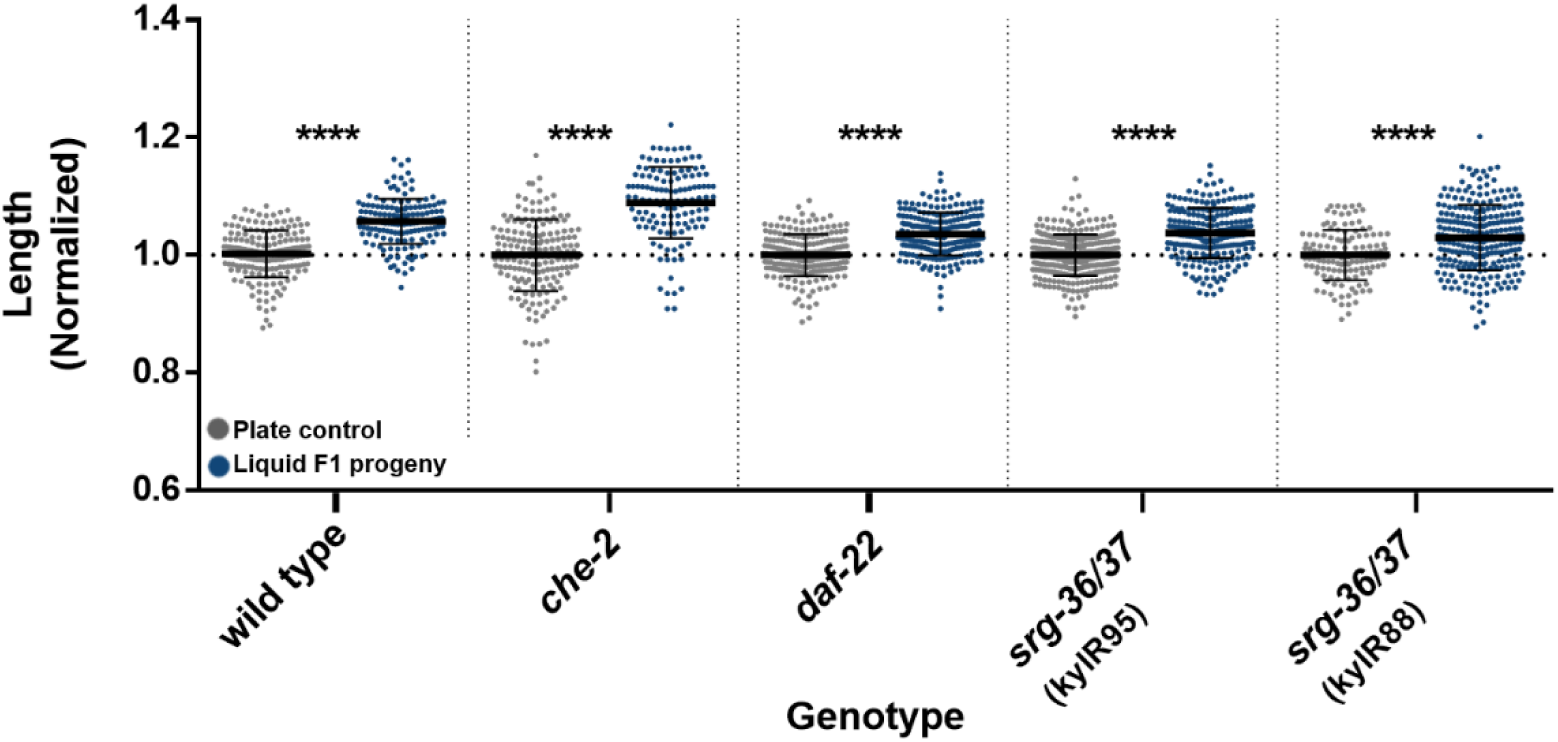
Progeny of liquid-grown worms are longer even in mutants defective in chemosensation and pheromone biosynthesis. The normalized lengths of progeny of liquid-grown mutants, defective in genes involved in dauer pheromone synthesis (*daf*-*22*), pheromone sensation (*srg*-*36/37*), stress response (*daf*-*16*), and chemosensation (*che*-*2*) are presented. N>100 per group. ^∗∗∗∗^p-value <0.0001, One-way ANOVA, Sidak’s multiple comparisons test. Error bars represent standard deviations.

CHE-2 is a protein with an unknown function that contains G-protein like WD-40 repeats. The sensory neurons’ cilia of *che*-*2* mutants are severely defective [39]. Accordingly, these mutants show defects in osmotic pressure avoidance [40], and are defective in chemotaxis to NaCl [39,40] and to many odorants [41]. We found that progeny of liquid-grown *che*-*2* mutants are also elongated, similarly to wild type worms. Therefore, sensation of secreted molecules does not appear to mediate the intergenerational response that we describe here (**Figure 3**).

DAF-16 is a transcription factor playing a pivotal role in the insulin/IGF-1 pathway, regulating stress responses, stress survival, longevity, fat metabolism, immunity and dauer formation [42]. DAF-16 was further implicated in inter-generational responses to stress [16]. Surprisingly, plate-grown progeny of liquid-grown *daf-16* mutants were elongated, similarly to wild type worms (**Figure 3**). This suggests that the DAF-16 dependent stress response is not involved in the liquid-induced heritable effect.

### Progeny of liquid-grown worms are longer even in mutants defective in inheritance of small RNAs and chromatin modifications

To examine whether the heritable changes in body length depend on inheritance of small RNAs or histone modifications, we examined mutants which are defective in production or inheritance of small RNAs and histone modifications. The nuclear argonautes HRDE-1 and NRDE-3 carry heritable small RNAs, and are required for the inheritance of dsRNA-induced silencing [43,44]. While NRDE-3 is expressed in somatic tissues and is important for inter-generational inheritance of somatic RNA interference [45], HRDE-1 is expressed in the germline and is required for the trans-generational inheritance (lasting multiple generations) of exogenous, dsRNA-derived small interfering RNAs (siRNAs) and endogenous siRNAs that target germline genes [43]. The amplified siRNAs that are carried by the NRDE-3 and HRDE-1 argonautes are produced by RNA dependent RNA polymerases (RdRP), such as RRF-1. RRF-1 is involved in the inheritance of piRNA-triggered endogenous siRNAs and viral-derived siRNAs [5,46]. We found that *hrde*-*1*, *nrde*-*3*, and *rrf*-*1* mutants, as well as additional mutants defective in siRNA biogenesis (*rde*-*1*, *rde*-*4*, *rrf*-*3* and *mut*-*16*) are not required for the inheritance of the elongated morphology (**Figure 4**). Previously, Histone-H3-Lysine-9 (H3K9) methylations were implicated in trans-generational inheritance of small RNAs [47–50]. H3K9 mono and di-methylations are deposited by the MET-2 methyltransferase [51], while H3K9me3 is deposited by the SET-25 and SET-32 methyltransferases [51,52]. We found that *met*-*2*, *set*-*25* and *set*-*32* mutants also inherit the elongated morphology from their liquid-grown parents (**Figure 4**). To conclude, the inter-generational inheritance of this phenotype does not appear to depend on the canonical epigenetic inheritance pathways that we examined.

**Figure 4.**
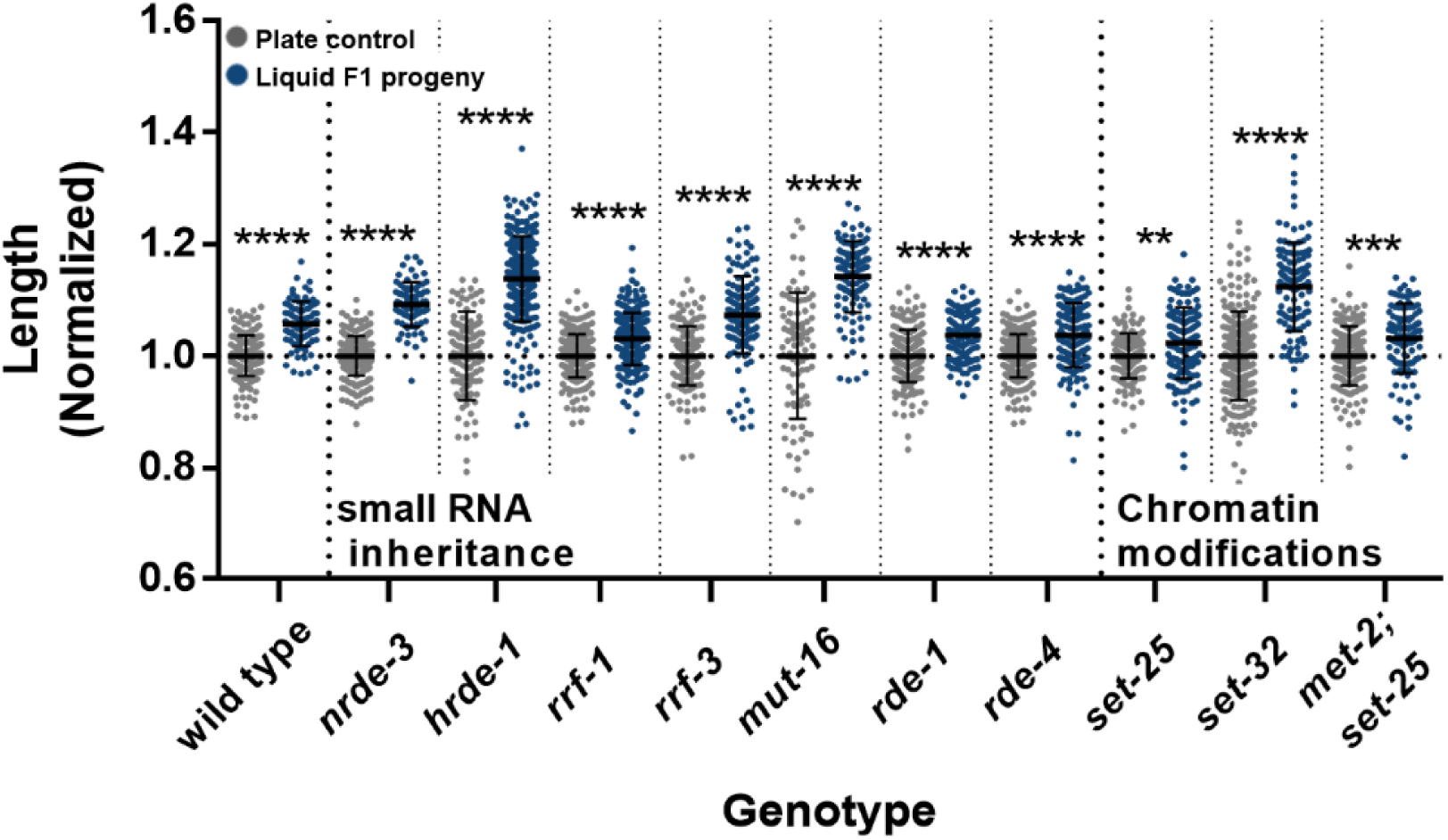
Mutants defective in inheritance of small RNAs and histone méthylations are also longer, if they derive from ancestors that were grown in liquid. The normalized lengths of progeny of liquid-grown worms mutant in genes involved in small-RNA biogenesis and inheritance (*hrde*-*1*,*nrde*-*3*, *rrf*-*1*, *rrf*-*3*, *rde*-*1*, *rde*-*4* and *mut*-*16* mutants) or chromatin methylation (*set*-*25*,*set*-*32 and met*-*2*;*set*-*25* mutants,) are presented. N>100 per group. ^∗∗∗∗^p-value <0.0001, ^∗∗∗^p-value=0.0001, ^∗∗^p-value=0.0084, One-way ANOVA, Sidak’s multiple comparisons test. Error bars represent standard deviations.

### Unlike the inter-generational effects, trans-generational inheritance of the morphological changes exhibits variability

Previously, certain environmental stresses, such as L1 starvation [53], starvation during dauer [11], and exposure to high temperatures [12], were shown to induce trans-generational effects that persist for more than two generations. To test whether the elongated body morphology can be inherited for more than one generation, we measured the body length of plate-grown worms that derive from liquid-grown parents in the F1, F2 and F3 generations. In contrast to the inter-generational changes to the worms’ size that were robustly inherited to the F1 generation, the trans-generational effect in the F2 and F3 generations was variable. While in some experiments we did observe trans-generational heritable changes, in other cases the effect was gone after 1-2 generations **(Figure 5).**

**Figure 5.**
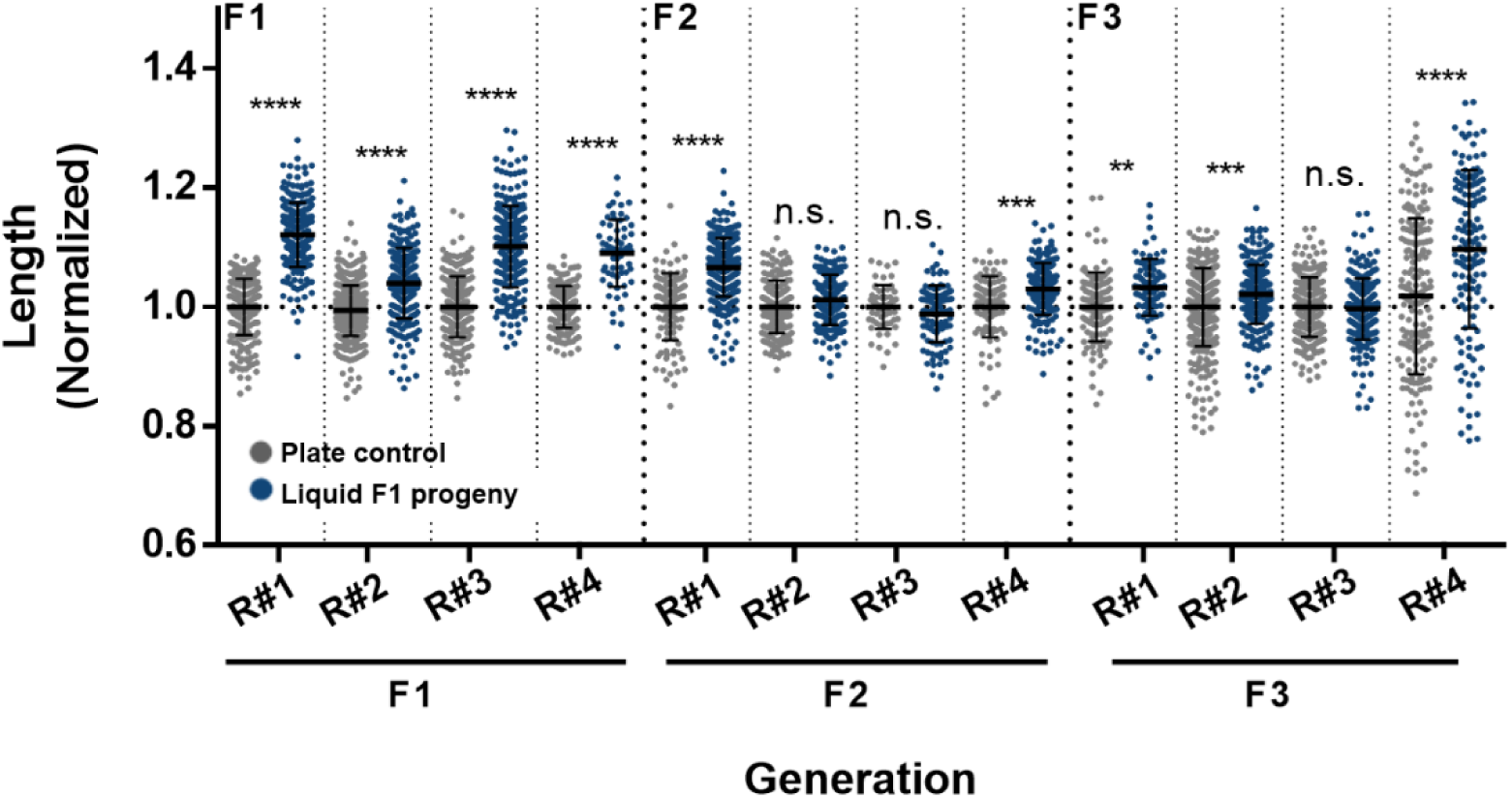
The liquid-induced longer morphology is not always inherited trans-generationally (in some experiments the effects peter out after 1 generation). The normalized lengths of progeny of liquid-grown worms and progeny of plate-grown controls along the F1, F2 and the F3 generations are presented. The measured length is normalized to the corresponding control worms of the matching generation and biological repeat. Data from four independent biological repeats are presented (N>150 worms per group). ^∗∗∗∗^p-value<0.0001, ^∗∗∗^p-value <0.005, ^∗∗^p-value=0.0015. One-way ANOVA, Sidak’s multiple comparisons test. Error bars represent standard deviations.

## Discussion

In this manuscript, we show that growth in liquid media under laboratory conditions changes the morphology of the worms and the morphology of their progeny. As briefly described above, there are rare reports of isolation of *C. elegans* from fresh water habitants [26]. There are also phenotypes, which are specific to worms that grow in liquid. For example, infection of *C. elegans* in liquid by the pathogenic bacteria *Leucobacter celer* was found to induce aggregation of worms (termed “worm stars”) [54]. Whether the morphological changes that we describe are adaptive in natural circumstances remains to be determined.

In addition to *C. elegans* other nematodes are routinely cultured in liquid cultures. Entomopathogenic nematodes such as *Steinernema* [21,22] and *Heterorhabditis* [23,24] are grown in liquid cultures as means of mass production of a biocontrol agent. However, liquid-grown entomopathogenic nematodes were found to have lower dauer stage recovery rates and pathogenicity [21,55,56], and much effort was invested in optimizing the different aspects of their mass production in liquid cultures [22]. Recently, a different nematode model organism, *Pristionchus pacificus*, was found to exhibit altered, narrow and elongated morphology when grown in liquid cultures [25]. Interestingly, the dimorphic mouth-form of *P. pacificus*, was found to respond to growth in liquid medium, however, the response was not heritable [25]. Several additional *Pristionchus* species, but not all, also showed mouth-form alternation in response to growth in liquid culture [25].

During the course of this study we could not identify mutants in which epigenetic pathways are effected, that are defective for the inter-generational inheritance of the elongated morphology. It is possible that such mutants do exist, and could be detected in more comprehensive screens. Alternatively, the inter-generational effects that we examined could transmit via many different molecular “agents” that do not necessarily affect epigenetics. Pheromone production and sensation, does not seem to be required for this effect, as mutants in the relevant pathways also exhibited the heritable liquid-induced effect. Changes in other molecules, such as metabolites or hormones, which are not necessarily self-propagating or “amplifiable” (in contrast to heritable small RNAs), could mediate such short term inter-generational inheritance as well.

Previously, environmental changes such as starvation and temperature shifts, both common in the lab, were shown to change the physiology of the worms and of its progeny. Together with growth in liquid, these manipulations should be taken into consideration when designing experiments with *C. elegans.* It is reasonable to assume that many other subtle characters are effected in the progeny of liquid-growth worms (for example behavior). To control for such changes, the worms’ growth conditions should be monitored for several generations before experimentation.

Inter-generational and trans-generational inheritance present a unique challenge: to compare between different experiments, the experimenter must keep track of the history of the culture, to make sure different experiments begin from the same starting point. There are currently no standard protocols for achieving this goal. We do not know yet how to “re-set” epigenetic inheritance. Previously, we used CRISPR/cas9 to deactivate HRDE-1, a factor which is important for small RNA inheritance, consequently erasing small RNA memory. This led to the re-setting of the accumulated sterility of chromatin mutants (Mrt phenotype) [49]. Since not all heritable effects are necessarily small RNA-mediated, other re-setting strategies should be devised in parallel.

The same principles that were discussed above with regard to inter- and trans-generational epigenetic inheritance in *C. elegans* could apply also to other organisms, and even to experiments conducted with cell lines. Several years ago, a controversy arose regarding the mislabeling of many cell lines [57]. While this is certainly a serious problem, it is worth noting that even when the correct cell line is used, cells in culture acquire mutations [58] (and differ in ploidy), and moreover non-genetic inheritance could create variability between cultures in different labs. Indeed, the “generation time” of the culture is a factor, as cells that were passaged many times can differ from recently thawed stocks [59,60]. RdRPs, which are crucial for small RNA inheritance in *C.elegans*, are not known to be conserved in humans [4], however, even if this mechanism is missing in other organisms, epigenetic memories could transmit to daughter cells via different feedback mechanisms [61,62]. For example, in human cells, genes known to be induced by IFN-γ were found to exhibit a more robust induction in cells that have previously experienced IFN-γ [63,64]. Importantly, this effect was maintained for 4–7 cell divisions, and was associated with H3K4 dimethylation [64]. Therefore, while the causative agents are not always characterized, it is clear that some information which is not encoded in the DNA sequence can persist for long durations of time. This type of inheritance should be taken into account in the experimental design.

## Methods

### Cultivation of worms

Standard culture techniques were used to maintain the nematodes on nematode growth medium (NGM) plates seeded with OP50 *Escherichia coli* bacteria. Extreme care was taken to avoid contamination or starvation, and contaminated plates were discarded from the analysis. For inheritance experiments worms were synchronized either by a standard Sodium hypochlorite bleaching techniques, or by letting adult worms lay eggs on OP50 seeded plates for two hours.

### Growth in liquid media

Worm eggs were seeded into flasks with 20 mL of M9 buffer (including 1 mM MgSO4, [18]) supplemented with 200 μl Penicillin-Streptomycin-Nystatin (Penicillin G Sodium Salt: 10,000 units/mL, Streptomycin Sulfate: 10 mg/mL, Nystatin 1,250 units/mL), 20 μl Cholesterol (5 mg/mL) and were fed with 200 microliter of 1 mL concentrated pellet of 50 mL OP51 *E. coli* bacteria that were previously grown over night (LB buffer + 100μg/mL Streptomycin). The liquid flask was shaken (120 RPM) at 20°C.

### L1 arrest experiments

Adult worms were washed using M9 buffer [18]. After removal of the buffer supernatant, ~100 μl pellet of worms was treated for 5 minutes with 1 mL of bleach solution (2 mL of 5% sodium hypochlorite, 7.5 mL double distilled water, 0.5 mL NaOH 10M). Next, 0.5 mL of M9 buffer was added and the solution was centrifuged (8000 RPM, 1 minutes). The resulting egg pellet was washed three times by removing the supernatant, adding fresh M9 buffer and centrifugation of 8000 RPM for 1 minutes to re-pellet the eggs. Around 400 eggs were left to hatch and arrest in the larval L1 stage on fresh plates for ~12 hours. Next, larval worms were transferred using M9 buffer to NGM plates seeded with OP50 bacteria. Worms were photographed 72 hours later.

### Strains used in this study

YY720: [*hrde*-*1* (tm1200) III.], YY158: [nrde-3(gg66) X.], RB798: [*rrf*-*1*(ok589) I.], GW638: [*set*-*25*(n5021) III.; *met*-*2*(n4256) III.], VC967: [*set*-*32*(ok1457) I.], MT17463: [*set*-*25*(n5021) III.], WM27: [*rde*-*1*(ne219) V.], WM49; [*rde*-*4*(ne301) III.], YY13: [*rrf*-3(mg373) II.], GR1823: [*mut*-*16*(mg461) I.], CF1038: [*daf*-*16*(mu86) I.]. CB1033: [*che*-*2*(e1033) X.], DR476: [*daf*-*22*(m130) II.]., EG6699: [*ttTi5605* II.; *unc*-*119(ed3)* III.], CX13249: [*srg*-*37*,*srg*-*38* (kyIR88) X.], CX13591: [*srg*-*36*,*srg*-*38* (kyIR95) X.]

### Worm length measurements

Worms were placed on empty NGM plates and photographed using ScopeTek DCM310 binocular camera. The worms’ length was measured using worm-sizer [65].

### Egg length measurements

Worms were allowed to lay eggs on an empty NGM plate, the laid eggs were photographed using ScopeTek DCM310 binocular camera. Egg length was measured using the particle analysis tool of the ImageJ software.

### Statistical Analysis

Two-Way ANOVA was used to compare the effect two categorical variables on the length of the worm’s progeny (this statistical analysis is relevant to the experiments shown on **Figures 1D, 2A, 2C, and Figures S2B and S2C**): A. biological condition (growth in liquid or on plates) and B. biological repeat (independent experiments conducted on separate groups of animals, on different days). One-Way ANOVA was used to compare the effect of growth in liquid medium on worm’s length between different genetic backgrounds (**Figures 3 and 4**), between different food regimes (**Figure 1C and Figure S1**), or between conditions across generations (**Figure 5)**. In cases of multiple comparisons between genotypes conditions, Sidak’s multiple comparison test was applied. Linear regression analysis and a statistical method equivalent to analysis of co-variance was used to compare the growth rates of progeny of liquid-grown worms and control worms throughout development (**Figure 2B and Figure S3**). Biological repeats are defined as independent experiments conducted on separate populations of animals, on different days. Statistical tests were performed using GraphPad Prism software (Graphpad Prism) version 6.

## Data accessibility

This article has no additional data.

## Author contributions

**I.L., R.B.** and **O.R.** conceived and designed the experiments; **I.L., R.B.**, **Y.L.** and **L.C.** performed the experiments; **I.L.**, and **O.R.** wrote the paper.

## Acknowledgments

We thank all the Rechavi lab members for the helpful comments and fruitful discussions. We thank Yoav Zeevi, Yoav Benjamini’s group, for his help with the statistical analysis. Some strains were provided by the CGC, which is funded by NIH Office of Research Infrastructure Programs (P40 OD010440). We thank the ERC (grant #335624), the Israel Science Foundation (grant #1339/17) and the Allen Discovery Center program through The Paul G. Allen Frontiers Group (12171). O.R. gratefully acknowledges the support of the Adelis foundation (#01430001000).

## Supplementary Figures

**Supplementary Figure 1.**
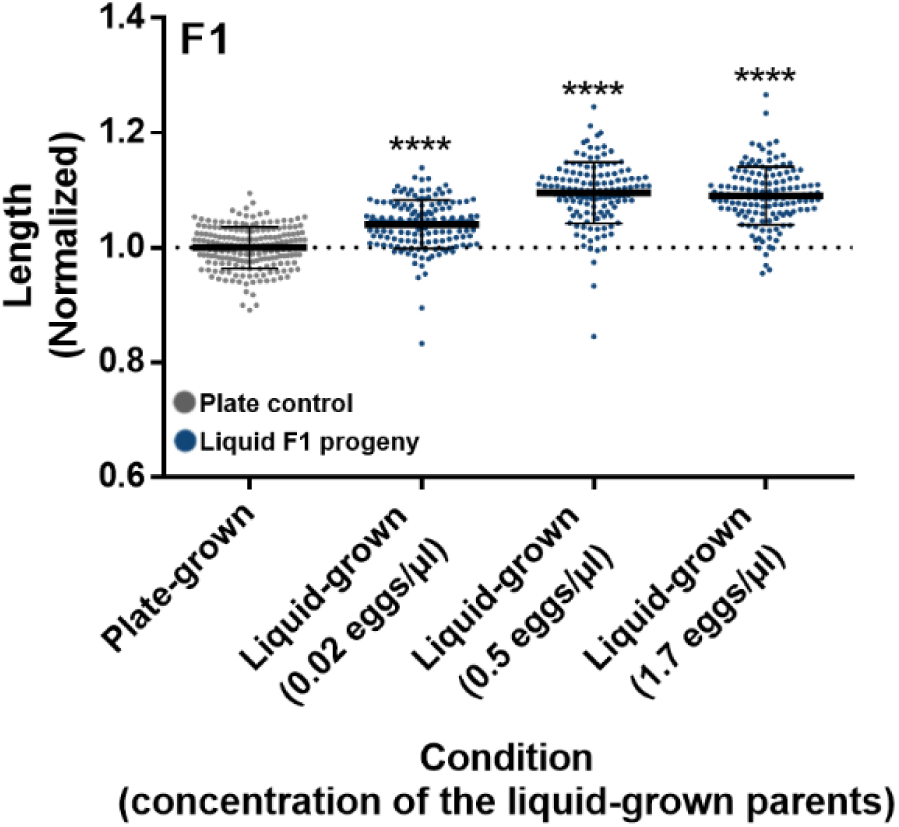
F1 progeny of liquid grown worms are longer irrespectively of the concentration in which their liquid-grown parents were grown. The normalized lengths of F1 progeny of worms grown in liquid in different concentrations (x-axis) and progeny of plate-grown controls are presented. The measured length is normalized to the corresponding control progeny of plate-grown worms. ^∗∗∗∗^p-value <0.0001. One-way ANOVA, Sidak’s multiple comparisons test. Error bars represent standard deviations.

**Supplementary Figure 2.**
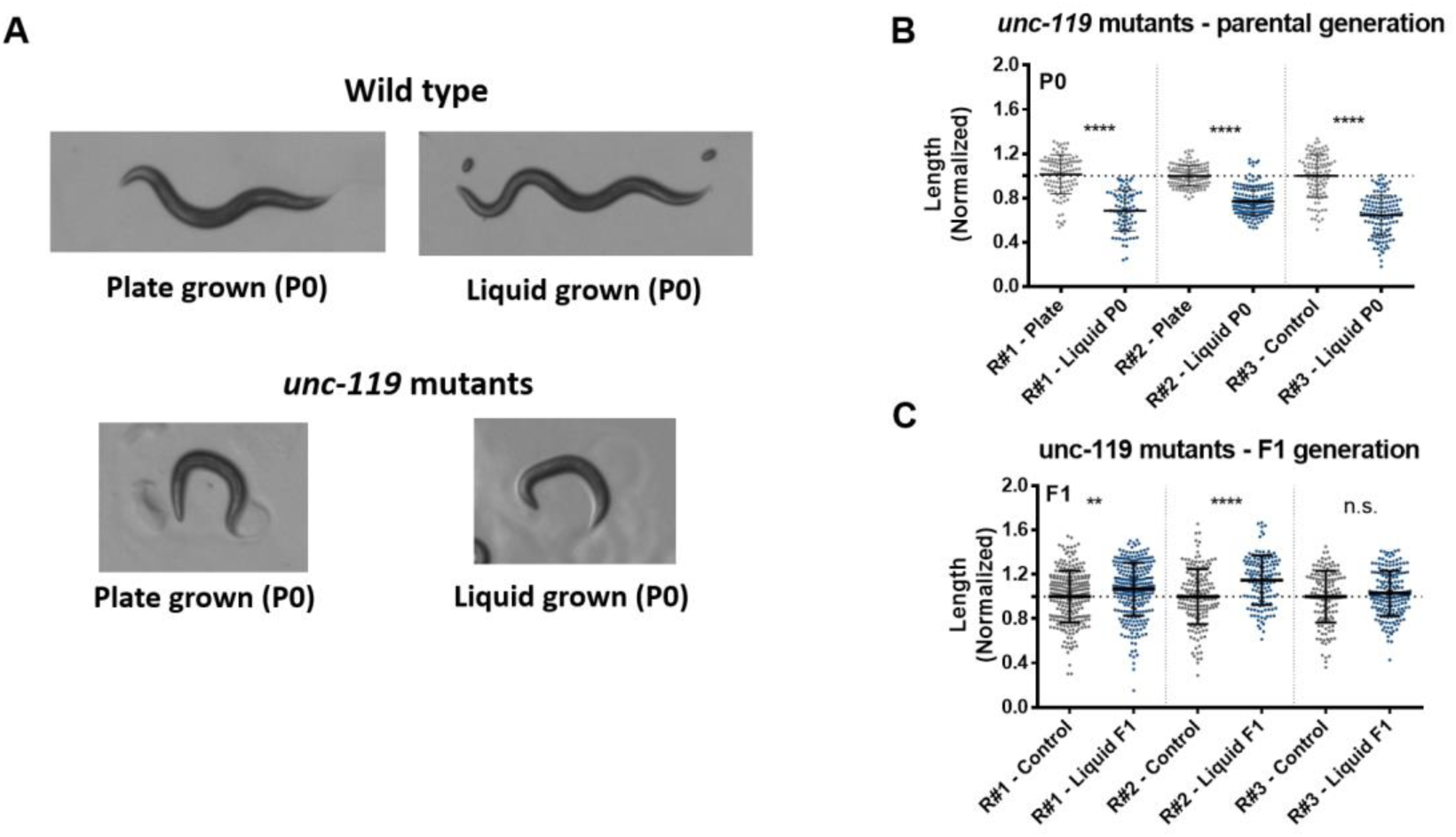
Immobile mutant worms do not alter their morphology when grown in liquid, but their progeny are longer. **(A)** Representative photographs of plate- and liquid-grown wild type and *unc*-*119* adult worms. **(B)** Quantification of the length of the parental unc-119 mutants grown either on plates or in liquid. The measured length was normalized to the corresponding control worms. Data from three independent biological repeats are presented (N>80). ^∗∗∗∗^p-value<0.0001, Sidak’s multiple comparisons test. **(C)** Plate-grown progeny of immobile *unc*-*119* mutant worms are longer than plate grown controls. The length of F1 plate-grown progeny of liquid-grown worms and progeny of plate-grown controls are shown. The measured length was normalized to the corresponding control worms. Data from three independent biological repeats are presented (N>130 worms per group). The p-value was determined using Two-way ANOVA with the two categorical variables: A. condition (growth in liquid or on plates) and B. biological repeat. The biological condition factor is significant (p-value<0.0001). The biological repeat factor and interaction effect are also significant (p=0.0024 for both). The biological condition factor explains more variance (2.76%) than the biological repeat or the interaction effect factors (1%). ^∗∗∗∗^p-value<0.0001, ^∗∗^p-value= 0.0027, Sidak’s multiple comparisons test.

**Supplementary Figure 3.**
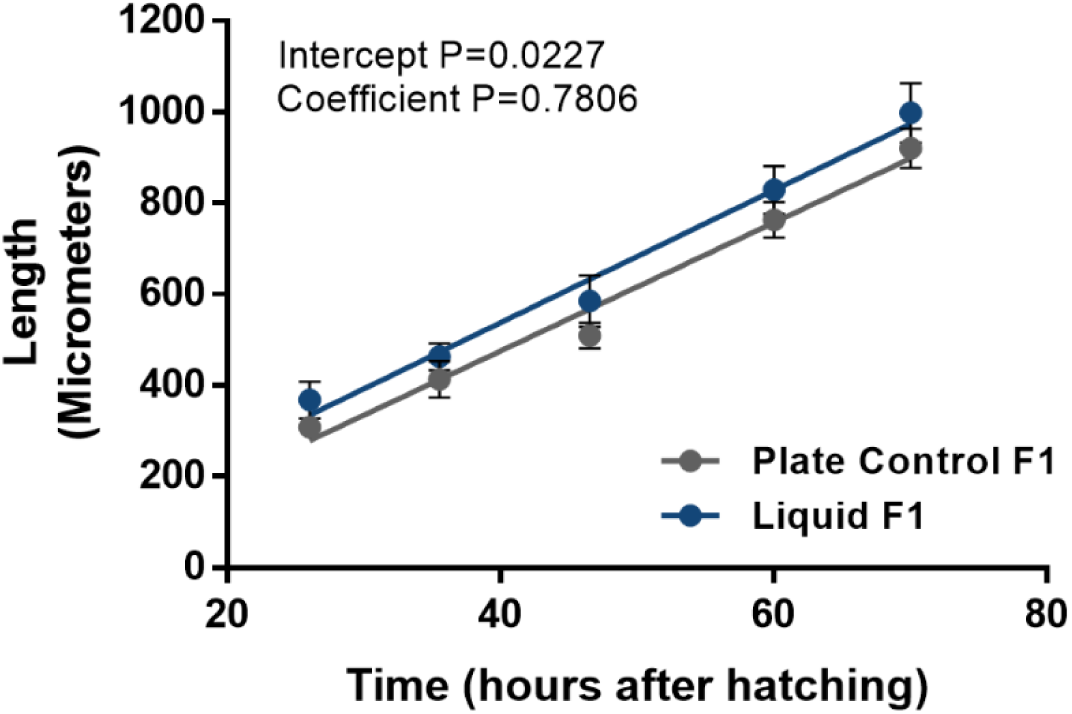
Linear regression analysis of growth rates of progeny of liquid-grown and control worms. Linear regression analysis of the length of progeny of liquid-grown worms and control worms at different time-points during the worm’s development. While the intercepts are significantly different (p=0.022), the rate of growth, or regression coefficients are identical (p=0.7806). Error bars represent standard deviations.

